# Statistical learning of bacterial growth in combinatorially constructed environments

**DOI:** 10.64898/2026.06.30.735472

**Authors:** Andrea Arrabal, Magdalena San Román, Juan Diaz-Colunga, Alvaro Sanchez

## Abstract

Optimizing growth conditions and culture media is a major goal in microbiology. A challenge is that nutrients can have complex, non-additive effects on growth. The fact that resources interact with one another has long been known in ecology, but systematic maps of resource interactions at all orders have been lacking and it is not known whether interactions are primarily low order or fundamentally complex. To tackle this problem, we have followed a full factorial design approach and measured the growth of seven different bacterial species in all possible combinations of 8 carbon sources under carbon-limiting conditions. Our approach allows us to directly estimate interactions at all orders. Even though all C-sources stimulate growth on their own as well as in combination with other nutrients, most of them can also have negative effects on growth when they are added to at least some nutrient mixtures. We show that the switch from positive to negative fitness effects is governed by global epistasis among resources. An analysis of variance shows that additive effects and pairwise interactions explain most of the variation in fitness, allowing us to train simple regression models that accurately predict bacterial growth in novel environments. The generality of these findings across seven different bacterial strains belonging to two different families indicates that interactions between carbon sources under carbon-limiting conditions may be generally learnable from a relatively sparse set of constructed environments, enabling the rational optimization of growth conditions.

## Introduction

Microorganisms live in chemically and nutritionally complex habitats, containing numerous different organic and inorganic resources. The collection of all these environmental resources modulate microbial growth and behavior, giving us a handle we could exploit to influence and manipulate microorganisms, from the very presence of bacterium in a microbiome (Beam et al., 2021; Walter et al., 2018) to its contribution to biotechnologically relevant functions (Eng & Borenstein, 2016; Sánchez et al., 2024). Fully exploiting this handle would require us to predict how targeted manipulations to the environment (e.g., adding and removing resources) will affect microbial growth.

This has been difficult to do due to the subtle context dependencies of molecular effects on microbial growth. The same environmental resource may have different growth and phenotypic effects in different environmental contexts and for different bacteria. When the effects of a resource on a bacterium are modified by the presence of other resources, these resources are said to interact (Tilman, 1980; Perrin et al., 2020; Sánchez et al., 2024). Both microbiologists and ecologists have long studied resource-resource interactions. A classic example is diauxic growth (Monod, 1942), where microorganisms will consume resources sequentially so that one resource will temporarily mask the effect of the other one. Other forms of interactions such as resource co-limitation have also been described in the ecology literature (Sperfeld et al., 2016; Held & Manhart, 2024; Held et al., 2024). Most past work on resource-resource interactions has focused on pairwise effects (Fannin et al., 1981; Harpole et al., 2011; Sperfeld et al., 2016), but pairwise interactions could also be in principle modified by the presence of additional resources, i.e. by higher-order effects. When we take these higher-order interactions into account, the number of all possible interactions between resources grows exponentially with their number: there are over one million potential interactions in an environment that contains as few as 20 different resources. Characterizing all of these interactions is impractical even for low-complexity habitats.

The problem of understanding how complex interactions between environmental resources (i.e. environment-by-environment interactions) shape bacterial growth yield is reminiscent of the similar problem of predicting how complex epistatic (i.e. gene-by-gene) interactions govern the fitness effects of genes in different genetic backgrounds (Bank, 2022). Both in genetics and in ecology, recent work has found that despite the potential for high-order epistatic interactions, additive effects and pairwise interactions often explain a large fraction of the variance in a fitness landscape (Khan et al., 2011; Skwara et al., 2023; Diaz-Colunga et al., 2024; Camacho-Mateu et al., 2026). We reasoned that the same may be true for resource-resource interactions. If this were generally the case for a wide range of bacteria and resources, it would suggest that optimizing growth media may be accomplished from a relatively small number of experiments.

To test this hypothesis, here we have generated all possible 255 combinations of eight carbon sources (i.e., a full factorial design (Diaz-Colunga et al., 2026)). We then independently cultured seven different bacterial strains (five from the Pseudomonadaceae family and two from the Enterobacteriaceae family) in each of the environments, and quantified their growth after 24 hours. We find that resources exhibit widespread interactions among them, and the fitness effect of a resource can be strongly affected by the presence of other resources. Most resources exhibited a dual behavior, switching between promoting or diminishing bacterial fitness when they were added to different background resource mixtures. This sign-variation in fitness effects can be explained, in part, by a phenomenon akin to that of global epistasis in genetics (Segrè & Marx, 2010; Kryazhimskiy et al., 2014; Otwinowski et al., 2018; Lyons et al., 2020; Bakerlee et al., 2022; Ardell et al., 2024), where resources act as growth-promoting nutrients in low fitness environments, but they have a negative effect on bacterial fitness when they are included in high-fitness environments.

These global epistasis patterns can be quantitatively explained from the underlying network of pairwise resource interactions. Although high-order resource-resource interactions can be detected for most species, they explain a relatively small fraction of the total variance in bacterial fitness across environments. In all seven strains tested, background-averaged additive effects and pairwise interactions amongst resources explained over 90% of the variance in bacterial growth. This low effective dimensionality allows us to train simple regression models to predict bacterial growth in novel resource mixtures, which we successfully implement for all seven species. Our results explain the past success of low-dimensional statistical models of nutrient-nutrient interactions and suggest that this may be a general feature of microbial growth media optimization.

## Results

### Bacterial resources act as either promoters or inhibitors of growth based on environmental context

As a model system to investigate how multiple environmental resources combine their effects to govern microbial growth, we compiled a set of seven bacterial strains (*Pseudomonas atacamensis, Pseudomonas putida* (P2), *Pseudomonas alloputida* (P3), *Pseudomonas aeruginosa* (PA), *Pseudomonas putida* KT2440-GFP (KT), *Salmonella enterica* y *Serratia marcescens*) and eight carbon sources including two monosaccharides (glucose and fructose), a complex sugar (soluble starch), a sugar alcohol (glycerol), and four organic acids (citrate, butyrate, acetate and ascorbic acid). As recommended in past studies in resource co-limitation (Sperfeld et al., 2016; Held et al., 2024), all resources were added at a low concentration (0.003 C-mol/L) to a M9 minimal medium (our base medium), ensuring that even when all eight carbon sources were added (to a final concentration of 0.024 C-mol/L), bacteria were still carbon-limited, a point we confirmed empirically (Supplementary Fig. 1). Most strains were able to grow on most of the resources as the sole carbon source (Fig. 1A; Supplementary Fig. 2); for instance, *P. atacamensis* grows on every resource except for ascorbic acid as the sole source of carbon (see Supplementary Fig. 2 for the remaining strains).

**Figure 1.**
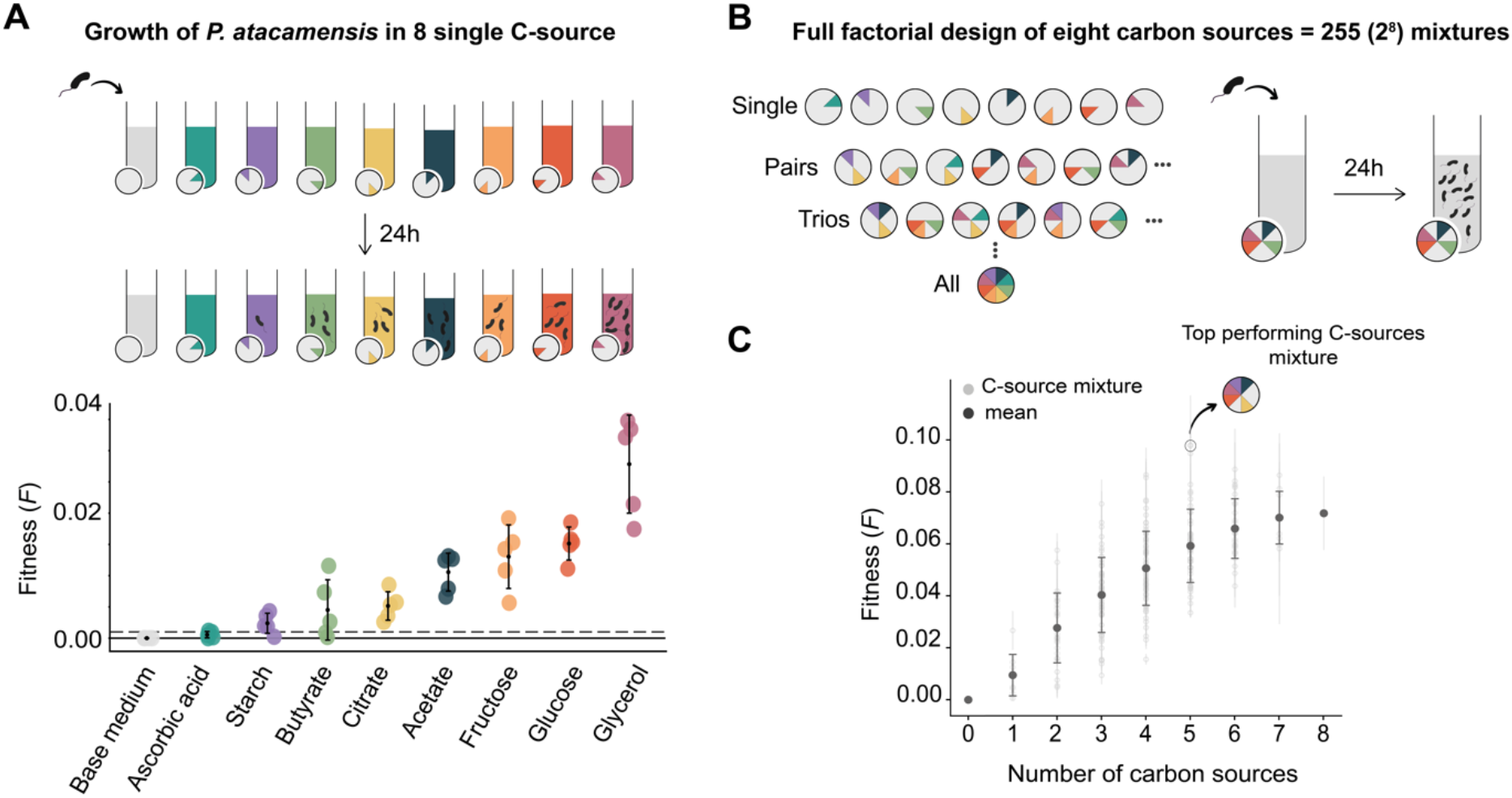
Effects of single carbon sources on the fitness landscape of Pseudomonas atacamensis. (A) We investigate the growth of Pseudomonas atacamensis on eight single carbon sources (acetate, ascorbic acid, butyrate, citrate, fructose, glucose, glycerol, and starch). Points represent biological replicate measurements and black error bars indicate mean ± standard deviation. All carbon sources except ascorbic acid support the growth of P. atacamensis as the sole source of carbon. Growth is defined as a mean OD_600_ above our limit of detection (0.001; dashed line), which coincides with the inoculation OD. (B) We constructed all 255 combinations of the 8 carbon sources. Each combination is represented by the color associated to each resource being present in the wheels. We then inoculated and grew P. atacamensis in each of the 255 environments. The growth yield was determined by the OD_600_ after 24 hours of growth. (C) We represent the growth yield as a metric of fitness of P. atacamensis in each of the environments, as a function of the number of resources in each environment. Light grey points represent fitness in each individual environment, and dark grey points and error bars indicate the average fitness (± standard deviation) as a function of the number of carbon sources present in the mixture. On average, we observed that increasing the number of resources results in increased fitness; however, the best-performing mixture (marked by the wheel containing the color-coded resources that are present therein) contains only five carbon sources.

To systematically investigate resource-resource interactions at all orders, we assembled every possible combination of the eight carbon sources, implementing a full factorial design consisting of 255 environments (2^8^ − 1 combinations; Fig. 1B) (Diaz-Colunga et al., 2026). Each environment contained 400 µL of base medium supplemented with the resource mixture, inoculated with 4 µL of overnight culture at an initial OD_600_ of 0.001. After 24 hours of incubation at each strain’s optimal temperature (Materials and Methods), we measured OD_600_ as a proxy for microbial reproductive fitness (*F*). Experiments were performed in five biological replicates on different days by the same experimentalist.

On average, we found that the fitness of all strains increased with the number of resources in the medium, as expected given the higher total carbon concentration (Fig. 1C; Supplementary Fig. 3). However, the optimal growth medium was generally not the one containing all carbon sources. For instance, for *P. atacamensis*, the highest fitness was achieved in a medium lacking fructose, ascorbic acid, and butyrate, despite the fact that this strain could grow in two of these resources as the sole C-source. This observation indicates that the effect of a resource may be contextual. Indeed, the marginal fitness effects for *P. atacamensis* ((*Δ*_*i*_*F*) = *F*(*b* + *i*) − *F*(*b*), where resource *i* is added to background mixture *b*) varied substantially across all backgrounds (Fig. 2). Although all resources had neutral or positive fitness effects on average, most of them exhibited negative fitness effects when added to at least a few background mixtures. For example, acetate addition to the growth medium reduced the fitness of *P. atacamensis* in 32/128 background mixture combinations. Although the specific distributions of fitness effects of all resources were different for each strain, the observation that adding a new carbon source to a background mixture (under carbon-limited conditions) could have negative fitness effects was widespread for all strains (Supplementary Fig. 4).

**Figure 2.**
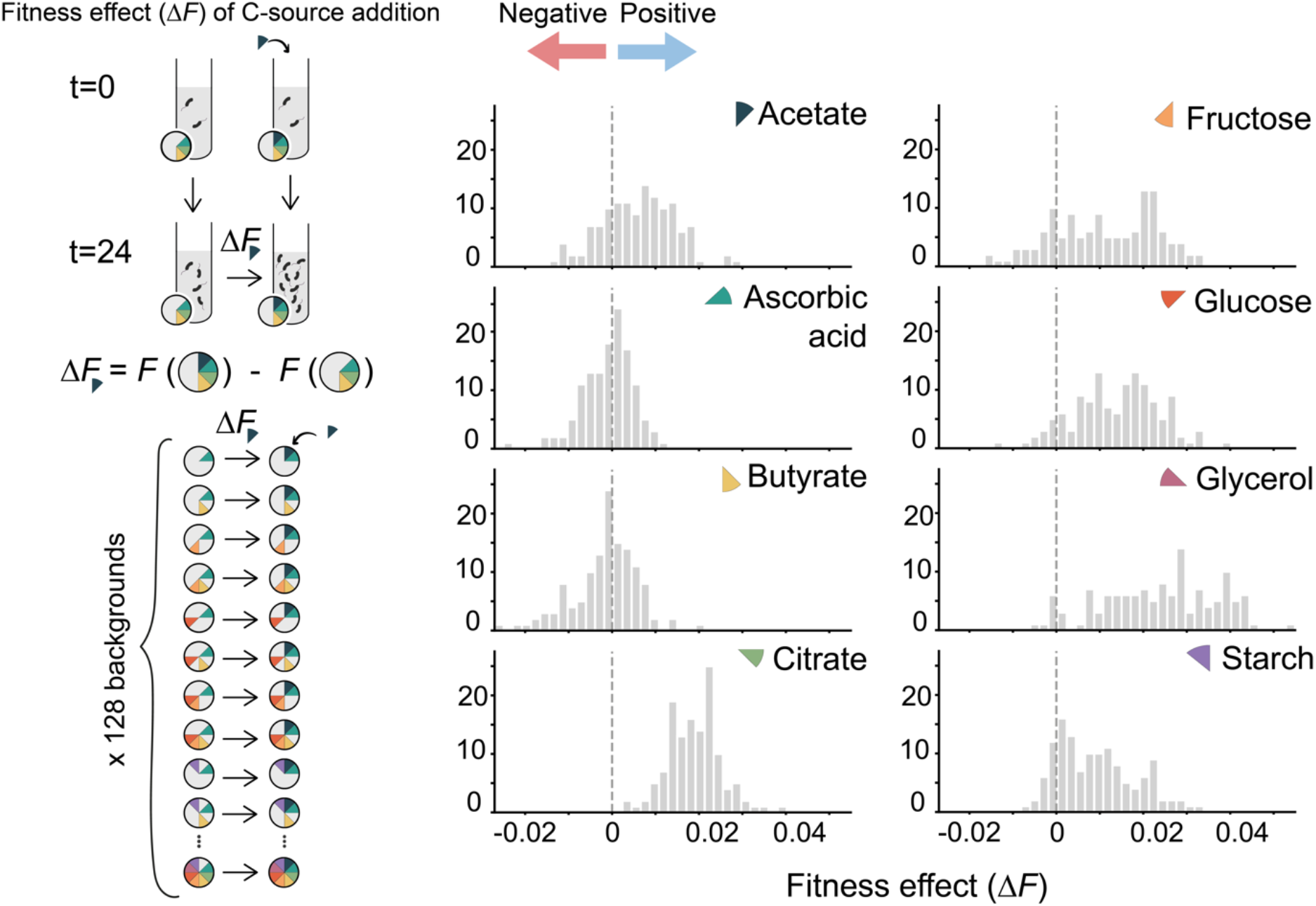
Context-dependent dual behavior of carbon sources. (Left) Experimental design showing how fitness effect (ΔF) measured by comparing the growth yield after 24 hours of growth in a mixture with and without a focal carbon source added, across 128 environmental backgrounds. (Right) Distribution of fitness effects for eight carbon sources (acetate, ascorbic acid, butyrate, citrate, fructose, glucose, glycerol, and starch, each represented by the same color code adopted in Fig. 1) across all 128 resource mixtures into which we can include them. Negative values (left of dashed line) indicate a detrimental effect on growth (growth suppression) while positive values (right) indicate fitness enhancement.

In sum, including a resource into a mixture can have substantially different fitness effects, both in magnitude and sign, depending on which other resources are part of that mixture. In genetics, the variance of the distribution of fitness effects of a new mutation reflects epistatic interactions (Reddy & Desai, 2021). It has been recently found that the aggregate effect of these interactions leads to the emergence of global epistasis trends, whereby part of the variation in fitness effect of a new mutation is explained by the fitness of the genetic background through simple, linear regression models (Reddy & Desai, 2021; Ardell et al., 2024; Diaz-Colunga et al., 2024). The slope, intercept and fraction of the variance explained by the regression can be mathematically connected to the statistical distribution of fitness effects and epistatic interactions. Therefore, we hypothesized that the same may be true for the fitness effect of resources. If this were indeed the case, and similar to what happens in genetics, it may be possible that the variation in sign effect of resources is driven by global epistasis, rather than a consequence of idiosyncratic interactions between specific sets of resources. We set out to test this hypothesis.

### The fitness effects of carbon sources exhibit patterns akin to global epistasis

To test whether widespread resource interactions lead to global epistasis patterns amongst resources, we re-examined the fitness effects data of *P. atacamensis* (Fig. 2). A manifestation of global epistasis is that part of the variation in fitness effect of adding resource *i* to different backgrounds can be explained by the fitness in the environmental background *b* to which *i* is added. To test this, we plotted the fitness effects of each resource *i* in each background environment *b* (*Δ*_*i*_*F*(*b*)) against the fitness of *P. atacamensis* in that same background environment (*F*(*b*)) (Fig. 3A). Consistent with our hypothesis, we found that a simple ordinary linear regression model of the form *Δ*_*i*_*F*(*b*) = *α*_*i*_ + *β*_*i*_*F*(*b*) (which we refer to as Fitness Effect Equations (FEEs) (Diaz-Colunga et al., 2024)) explained a fraction of the variance ranging from 23% for starch to 51% for fructose (Fig. 3B). A notable exception was citrate (*R*^2^ = 0.02, slope ≈ 0), whose fitness effects were largely independent of background fitness. This exception proved instructive: as we show below, both the widespread predictability observed for most resources and this anomaly can be fully explained by the underlying network of pairwise resource interactions

**Figure 3.**
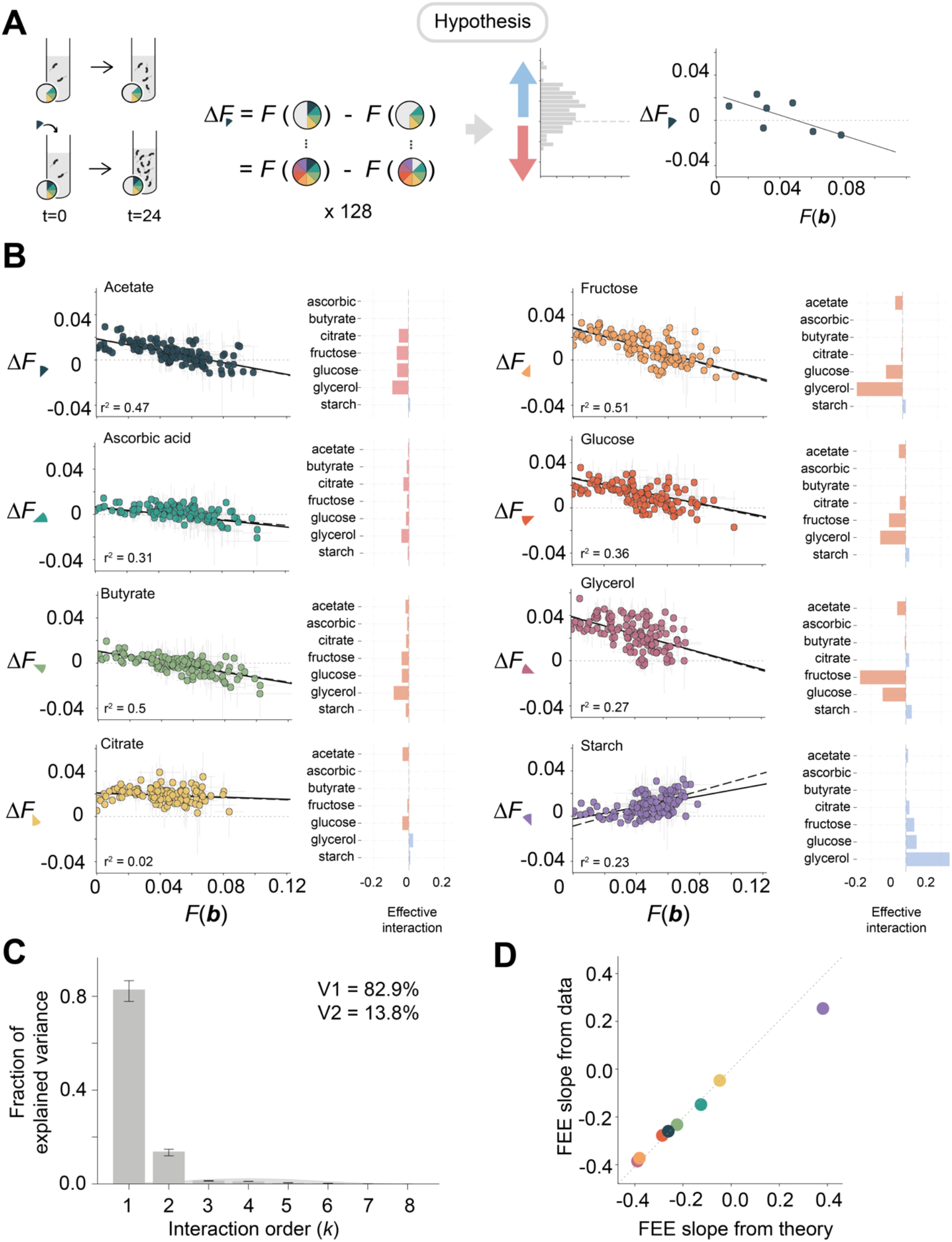
Environmental parallel to global epistasis. (A) We test the hypothesis that a phenomenon akin to global epistasis amongst mutations might explain the fitness effects of including of adding a focal resource to a background environment. To this end, we will test whether the fitness effect of a new resource (Δ.F) can be predicted from an ordinary linear regression on the fitness of the background environment (F(b). (B) We plot Δ.F versus F(b) for P. atacamensis across all 8 carbon sources. Solid lines show fitted regression lines; dashed lines indicate predicted FEEs from pairwise resource–resource interactions and individual fitness effects of non-focal resources (see Materials and Methods). Right panels show the sign and magnitude of effective interactions between the added carbon source and the other resources. (C) Fraction of total explained variance in fitness for P. atacamensis across the 255 environments attributable to interactions at all orders. Additive effects and pairwise interactions explain > 96% of fitness variation across all environments. (D) Observed FEE slopes for P. atacamensis obtained from the linear regressions versus the expected slopes predicted from the pairwise interaction model (Eq. 1). The tight correlation validates that resource interactions decompose into predictable pairwise effects, explaining the global epistasis pattern.

To understand why global epistasis emerges from resource interactions, we resorted to the theory of fitness landscapes in quantitative genetics, which provides a principled framework for decomposing how epistatic terms combine to give rise to the slope of the FEEs. Applied to interactions between resources, this theory establishes that the slope of the FEE regression (*β*_*i*_) is determined by the pairwise epistatic interactions between resource *i* and all other resources in the background resource mixture (Reddy & Desai, 2021; Diaz-Colunga et al., 2023a; Diaz-Colunga et al., 2023b):

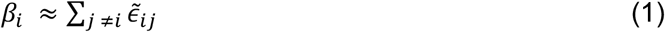

In the above equation, the effective pairwise interaction 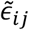 between resource *i* and *j* is defined as:

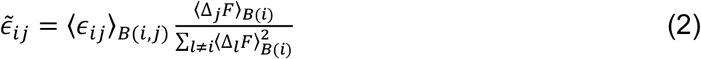

where *⟨*ϵ_*ij*_*⟩*_*B*(*i,j*)_ is the pairwise epistasis between resource *i* and *j*, averaged across all background mixtures *B*(*i, j*) in which neither resource is present; *⟨Δ*_*j*_*F⟩*_*B*(*i*)_ represents the average fitness effect of resource *j* over all backgrounds that do not contain resource *i* (*B*(*i*)). Eq. 2 is derived in the limit where high-order interactions are negligible (Reddy & Desai, 2021; Diaz-Colunga et al., 2023a; Diaz-Colunga et al., 2023b), a limit that, as we show in Fig. 3C and discuss later, is met in our experiments.

Equations 1-2 quantitatively connect the slopes of the FEEs for all resources to the pairwise resource-resource interactions involving our focal resource *i* and the individual fitness effects of the non-focal resources *j*. All of these parameters can be independently determined from our data (Materials and Methods), and we can therefore use the above equations to estimate the expected regression slope from the epistatic interactions amongst resources. As shown by the dashed lines in Fig. 3B, The FEEs predicted from the interactions for *P. atacamensis* match very well the actual regression lines (solid lines), and the observed regression slopes are in excellent agreement with the theoretical predictions from Eq.1 (Fig. 3D). This finding was general and consistent for all other six strains as well, and for all resources (Supplementary Fig. S5 and Supplementary Fig. S6).

Because Equations 1 and 2 successfully predict the regression slopes, we can also use them to interpret the interaction mechanisms underlying the FEEs. For instance, the steep negative slope observed for acetate can be explained by its strong negative effective pairwise interactions with citrate, fructose, glucose and glycerol (Fig. 3B, right panels), which dominate over the negligible interactions that acetate has with ascorbic acid, butyrate, and starch. In contrast, the positive slope of starch is driven by positive pairwise interactions it has with citrate, fructose, glucose and glycerol (Fig. 3B, right panels). Finally, the near-flat FEE observed for citrate reflects the weaker interactions it has with the other resources, which also include both negative and positive interactions that partially cancel each other out. The same analysis was conducted for all other strains as well, finding qualitatively similar results (Supplementary Fig. S5).

The fact that additive and pairwise effective interactions explain the observed patterns of global epistasis so well suggests that high-order interactions have a relatively small contribution to the fitness effects of new resources under our conditions. Indeed, as we discussed above, additive effects and pairwise interactions explain over 96.7% of the variation in fitness across all environments for *P. atacamensis* (Figure 3C). This result was quantitatively and qualitatively similar for the other species as well, leading us to conclude that higher-than-pairwise interactions have a very small effect on the fitness effects of new resources under our conditions (Supplementary Fig. 7).

Given the overwhelming dominance of low-order interactions and the tiny contribution of higher-order effects, we reasoned that statistical models including additive and pairwise interactions only will be capable of predicting bacterial fitness in new environments, even when trained on relatively small-sized and sparsely sampled training sets. We set out to test this hypothesis.

### Additive effects and pairwise interactions predict growth in out-of-sample environments

To test our hypothesis, we developed a statistical framework to predict bacterial fitness in complex, combinatorially constructed environments. We modeled the growth yield *F*(*x*) as a function of resource presence/absence using linear models that contained up to pairwise interactions:

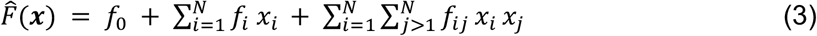

Where *x*_*i*_ is a binary variable that may take values {+1/-1} depending on whether species *i* is present or absent in the mixture, respectively. The fitting parameters *f*_*0*_, *f*_*i*_, *f*_*ij*_ reflect the average fitness, (background averaged) additive effects of resource *i*, and (background averaged) interaction between resources *i, j*, respectively. To test the predictive power of this model, we randomly partitioned the full factorial set of 256 environments (including the null, which contained no carbon) varying the fraction of observations in the training set from 5% (12 environments) to 90% (230 environments). We fitted Eq. 3 to the training set using Lasso or Ridge regularization via inner-fold cross-validation to avoid overfitting (Skwara et al., 2023). The trained models were then used to predict fitness for the held-out test set environments. Model performance was quantified using:

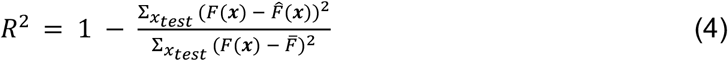

where *F*(*x*) is the observed growth yield for resource mixture 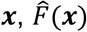 is the model prediction, 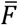 is the mean fitness across the test set, and the sums are computed over all assemblages in the test set.

We first tested the predictive power on the model for *P. atacamensis* (Fig. 4A). The results were strongly encouraging. Across 100 independent random partitions at each training fraction, the median *R*^2^ for the linear model was 0.86 at 17% training fraction, increasing to *R*^2^ > 0.93 at 37% and *R*^2^ > 0.94 at 70% or higher (Fig. 4B). We then repeated this analysis for the other six species with quantitatively and qualitatively similar results (Supplementary Fig. S8). An alternative statistical modeling strategy that leverages global epistasis instead (the stitching method described in Diaz-Colunga et al., (2024) (Supplementary Fig. S9)) produces similar results.

**Figure 4.**
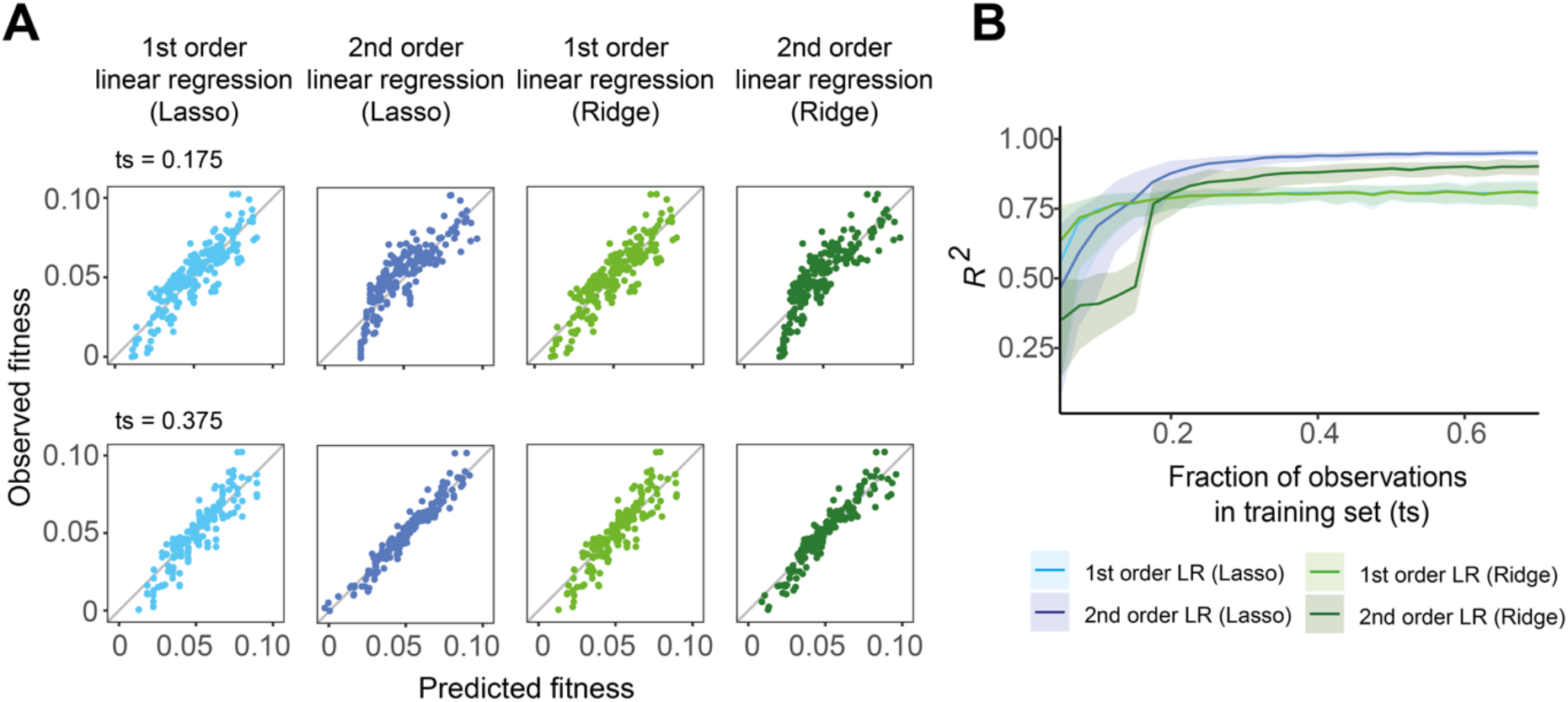
Linear regression models including up to pairwise interactions accurately predict microbial growth out of sample. (A) Observed versus predicted fitness values for first-order (1st order) and second-order (2nd order) linear regression models with Lasso and Ridge regularization at training set (ts) sizes (fraction of all resource mixtures) of 0.175 and 0.375 in P. atacamensis. (B) Relationship between training set size and model prediction accuracy (R^2^). Lines represent median R^2^ across 100 independent random partitions. Second-order models consistently outperform first-order models above ts=0.2, with differences becoming more pronounced as training set size increases.

In conclusion, our results indicate that additive effects and pairwise resource interactions are sufficient to capture microbial growth, and that a sparsely sampled training set of environments is generally sufficient to predict microbial growth in new, combinatorially constructed resource mixtures. The fact that this was true for all seven strains, belonging to two different families, suggests that at least under carbon limited environments our conclusions hold some generality.

## Discussion

We have systematically examined the patterns of nutrient interactions among eight bacterial resources for five different strains of Pseudomonas and two strains of Enterobacteriaceae. All resources in our study are carbon sources, and our experiments were carried out under carbon limiting conditions, where the response of bacteria to increasing concentrations of carbon is still linear, on average, and the existence of interactions is not granted. Yet, while additive effects capture a large proportion of the variance in resource effects for all species, pairwise interactions introduce context dependence, such that the effect of a given resource on bacterial growth depends on which other resources are also available.

Remarkably, we find that most resources have negative effects on the fitness of individual strains when they are included in at least some resource mixtures. This is true even when said resources can be utilized for growth as the sole carbon source by those strains, and their fitness effect is on average positive across all background resource mixtures. This sign-reversal effect is generally well explained by negative interactions between resources, whereby a resource promotes growth alone but can become detrimental for growth in the presence of other resources.

While the existence of resource interactions might appear to complicate the task of predicting the relationship between growth medium composition, we find that additive effects and pairwise interactions dominate, and high-order interactions contribute very little to microbial growth. Furthermore, a parallel phenomenon to global epistasis also emerges for resource interactions and simple linear regression models may explain the fitness effects of adding a new resource into a complex resource mixture as a function of the growth yield achieved in the complex nutrient mixture itself. We show that these linear models can be also exploited to accurately predict microbial growth in newly constructed environments, and thus represent a promising tool to optimize culture conditions in microbiology.

The fact that nutrients may combine non-additively to determine bacterial growth has been known for a very long time (Monod, 1942). Yet, the systematic empirical characterization of high-order nutrient combinations has been rare, and most mechanistic models of microbial consumer-resource interactions assume additive nutrient effects (Goldford et al., 2018; Zhang & Becks, 2025; Moran et al., 2026; Corral López et al., 2026). Other mechanistic approaches, such as those based on genome-scale metabolic models (Dukovski et al., 2021; Takano et al., 2023), tend to assume that growth is optimal, implicitly biasing nutrient interactions to be positive or neutral (with the possibility of diauxic-based negative magnitude being incorporated through additional cost-related constraints (Mori et al., 2016)).

While the possibility of antagonistic sign-effect nutrient interactions (those that alter the sign effect of a nutrient, making it detrimental) has been contemplated by theoretical models, and such interactions have been reported in plants and fungi (Fageria, 2001; Rietra et al., 2017; Deveau et al., 2018; Mishra et al., 2024), the extent to which antagonistic interactions between nutrients are realized in bacteria had not been studied systematically. Our results suggest that future theoretical work would have to consider them if we wish to investigate questions as pressing as how biodiversity changes with growing nutrient diversity, or how nutrient exchanges may modulate species coexistence.

Our observation that a resource can act as either a growth promoter or inhibitor depending on the environmental background complicates the design of optimal environments for microbial processes, posing significant challenges with clear biotechnological implications. Moreover, this dual behavior of resources has far-reaching consequences for ecology and evolution. For instance, we tend to view resource exchange as producing facilitation and favoring coexistence (Pande et al., 2015; Rappaport et al., 2025; Sulheim et al., 2026); yet, antagonistic interactions may cause cross-feeding to be inhibitory instead. This should lead us to reconsider the relative effect that cross-feeding may have on microbial interactions and microbial coexistence (D’Souza et al., 2018; Goldford et al., 2018; Kost et al., 2023).

While our findings provide what we hope are valuable insights into resource interactions and their effects on microbial fitness, several limitations should be acknowledged. Our analysis was based on only eight resources, all of which are carbon sources and each tested at a fixed concentration under carbon-limiting conditions. Therefore, our experimental design may not be capturing the full range of potential resource interactions that are likely to occur at higher concentrations, including diauxic growth. Additionally, our experiments were conducted with a limited set of seven strains of Gammaproteobacteria, and thus our study is inevitably limited phylogenetically. Further investigations involving a broader diversity of resources, concentrations, and microbial taxa will be necessary to fully understand the generality and scope of these interactions. Notwithstanding these limitations, the methodology we have developed here makes it possible to rapidly and inexpensively implement a full factorial design of additional resources, growth media and culture conditions, and analyze the resulting data using the framework of epistasis, which we borrowed from genetics. By incorporating a greater diversity of resources, varying concentrations, and broader microbial taxa, such studies will deepen our understanding of how resources combine their effects and reveal their ecological and evolutionary consequences.

## Supporting information

Supplementary Information (Suppl. Figures and Table from Statistical learning of bacterial growth in constructed environments).

## Acknowledgements

We are especially grateful to M.D. Fernández-de-Bobadilla for their help with the WGS analyses. We also thank all members of the Sánchez lab for helpful discussions. We thank Sara Hernando-Amado, Esteban Martínez, and Francisco Rey for kindly providing several strains used in this work.

## Funding

This work has been funded by the European Union (ERC, ECOPROSPECTOR, 101088469). We also thank Adisseo France S.A.S for funding the initial stages of this project. M.S.R acknowledges support from the Juan de la Cierva programme (JDC2021-046960-I). J.D.-C. acknowledges support from grant RYC2023-045580-I funded by MICIU/AEI/10.13039/501100011033 and by the “European Union NextGenerationEU/PRTR”, and from RyC-MaX-CSIC grant 20252MAX002 funded by the Spanish National Research Council (CSIC). The views and opinions expressed here are, however, those of the author(s) only and do not necessarily reflect those of the European Union or the European Research Council. Neither the European Union nor the granting authority can be held responsible for them.

## Competing interests

This work received partial financial support from Adisseo France S.A.S through a research collaboration agreement. The authors declare no competing interest.

## Materials and Methods

### 1. Bacterial strains and culture conditions

Bacterial strains used in this work were *Pseudomonas atacamensis, Pseudomonas putida* (P2), *Pseudomonas alloputida* (P3), *Pseudomonas aeruginosa* PA14 (PA) (Hernando-Amado et al., 2022), *Pseudomonas putida* KT2440-GFP (KT) (Koch et al., 2001), *Salmonella enterica* and *Serratia marcescens*. Strains *P. atacamensis*, P2 and P3 were isolated from different geographical locations in Madrid, Spain. Environmental samples were collected with sterile tweezers into 50 mL sterile tubes. One gram of each sample was allowed to sit at room temperature in distilled water. After 48 hours, 10 μL of each sample was inoculated onto minimal growth media supplemented with 0.2% glucose plates. A single colony from each strain was resuspended in liquid M9 medium with 0.02% glucose and incubated at corresponding temperature. Environmental samples were taxonomically classified based on whole-genome sequencing (WGS). Strains *P. atacamensis*, P2 and *S. marcescens* were incubated at 28°C whereas P3, PA, KT and *S. enterica* were grown at 37°C. Cell stocks were stored at −80°C in 40% glycerol.

To establish a strictly carbon-limited growth regime, preliminary calibration experiments were performed using a defined minimal medium (M9; hereafter referred to as base medium) composed of Na_2_HPO_4_ ·2H_2_O (48 mM), KH_2_PO_4_ (22 mM), NaCl (9 mM), and NH_4_Cl (19 mM), supplemented with MgSO_4_ (2 mM) and CaCl_2_ (0.1 mM). Cultures grown in Luria-Bertani (LB) medium were harvested and washed three times with sterile phosphate-buffered saline (PBS; 137 mM NaCl, 2.7 mM KCl, 10 mM Na_2_HPO_4_ ·2H_2_O, and 1.8 mM KH_2_PO_4_) to eliminate nutrient carryover. Cell suspensions were inoculated at 1% (v/v) into 96-well microplates containing 200 μL of fresh base medium supplemented with glucose. A 16-step two-fold serial dilution series was prepared starting from 1.6% (w/v) glucose. Growth was monitored for 24 hours by measuring optical density at 600 nm (OD_600_) using a Multiskan SkyHigh microplate reader (Thermo Fisher Scientific). Based on the resulting calibration assays, an individual resource concentration of 0.003 C-mol/L was selected. This concentration ensured that even in the maximum-complexity mixtures containing all eight carbon sources, the cumulative total carbon concentration would not exceed 0.024 C-mol/L, which remained within the linear carbon-limited growth regime for all strains tested.

### 2. Combinatorial assembly experiment and biomass quantification

Resources mixtures were assembled from eight carbon sources, including two monosaccharides (glucose and fructose), one complex sugar (soluble starch), one sugar alcohol (glycerol), and four organic acids (sodium citrate, sodium butyrate, acetate, and ascorbic acid) (Details in Table 1). Carbon source stock solutions were prepared at 8× concentration so that, after the combinatorial assembly protocol, each individual carbon source was present at a final concentration of 0.003 C-mol/L. Carbon source stock solutions were sterilized using 0.22 μm filters (Fisherbrand).

To prepare the starting inoculum, bacterial cells from a glycerol stock were streaked onto LB agar plates (Condalab), and incubated overnight at their respective strain-specific temperatures. A single colony was inoculated into a 50 mL conical tube containing 25 mL of LB broth (Condalab) and grown overnight under static conditions at the strain-specific temperature. Cells were harvested by centrifugation at 2500 rpm (1215 x *g*, Thermo Scientific SL Plus Centrifuge series) for 20 min. Cell pellets were washed three times with PBS, repeating the centrifugation step after each wash. Cultures were resuspended in base medium and diluted to an optical density at 600 nm (OD_600_) of 0.1.

Following a previously described protocol (Diaz-Colunga et al., 2026), we assembled the full combinatorial landscape of carbon sources by generating 256 unique combinations of 8 carbon sources in 96-deep well plates (Astik). Each well contained 400 μL of medium and was inoculated with 4 μL of the starter culture corresponding to the strain tested (1:100 dilution). Cultures were incubated for 24 hours at strain-specific temperatures. After incubation, plates were homogenized by automated pipette mixing using a MINI 96 electronic pipette (Integra Biosciences) prior to biomass quantification. Thirty-two negative control wells containing fresh base medium, both with and without carbon sources, were included per plate configuration. The entire combinatorial landscape was assayed across five independent biological replicates for strains *P*.*atacamensis*, P2, and P3; four replicates for KT, *S*.*enterica*, and *S*.*marcescens*; and three replicates for PA.

### 3. Quantification of fitness

The growth yield from an inoculation density of 0.001 was measured as the change in optical density at 600 nm over a 24-hour growth period. This was our metric of fitness throughout this paper. OD_600_ measurements at 24 hours were background-corrected by subtracting the OD_600_ of a reference well containing the base medium inoculated with the organism but without any carbon source, to account for background signal and optical density derived from the inoculum. Experimental data were corrected for batch effects and measurement noise using a Bayesian generative model that jointly estimates the underlying biological signal and systematic inter-replicate variation, following the methodology described in Camacho-Mateu et al., (2026).

### 4. Fitness effect calculation

To quantify the context-dependency of carbon sources effects on microbial fitness, we measured the marginal fitness contribution of each individual resource *i* (Δ_*i*_*F*) across different background resources mixtures. For each background mixture lacking resource *i*, the fitness effect was calculated as Δ_*i*_*F* = *F*(*b* + *i*) − *F*(*b*), where *F*(*b*) represents the microbial fitness of the background mixture, and *F*(*b* + *i*) represents the fitness of the identical background supplemented with resource *i*. Microbial fitness was defined as the endpoint biomass yield, quantified as optical density at OD_600_ after 24 hours of growth under the corresponding experimental conditions. For each resource *i*, Δ_*i*_*F* values were computed across all background combinations in which the resource was absent, generating a distribution of context-dependent fitness effects.

### 5. Quantification of functional effective interactions

The methodology for quantifying functional effective interactions was adapted from Díaz-Colunga et al., (2024). For each strain, the effective interaction between resources *i* and *j* (ϵ_*ij*_) was quantified as explained in the main text:

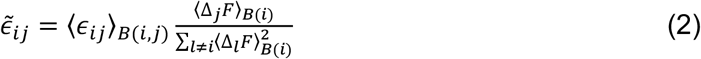

The term ϵ_*ij*_ represents the deviation between the fitness of background mixtures that contains both resources *i* and *j* with respect to the additive expectation that they do not interact, that is:

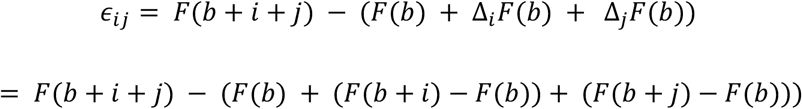

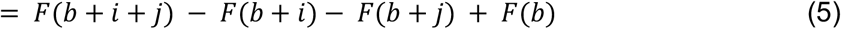

To quantify the average *⟨*ϵ_*ij*_*⟩*_*B*(*i,j*)_ as it appears in Eq. 2, we applied Eq. 5 to every possible background mixture *b* not contain resource *i* nor *j*, and took the average over all backgrounds.

### 6. Variance Decomposition via Walsh-Hadamard Expansion

To quantify the relative contribution of resources interactions at different orders to the total fitness variance of the landscape, we employed Walsh-Hadamard basis decomposition (Poelwijk et al., 2016; Skwara et al., 2023). The fitness *F*(*x*), defined over all possible resource configurations *x* ∈ {0,1} ^N^, is expanded as:

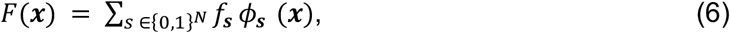

where *Φ*_*s*_(*x*) = (−1)^*s*(*x*+1)^ are the Walsh-Hadamard basis functions and *f*_*s*_ are the Walsh-Hadamard coefficients obtained using the uniform inner product over {0,1} ^N^:

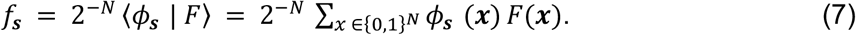

This decomposition provides an exact orthogonal decomposition of the total fitness variance across all mixture configurations, defined as:

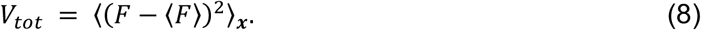

The order resolved variance involving exactly *n* resources is given by:

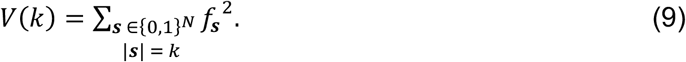

where |*s*| denotes the order of interaction (number of resources involved). This formulation is exact only when the all configurations are observed; otherwise orthogonality is lost (Camacho-Mateu et al., 2026). Eq. 9 allows us to determine which orders of resource interactions (individual effects, pairwise interactions, higher-order effects, etc.) contribute most substantially to the overall fitness heterogeneity of the landscape (Camacho-Mateu et al., 2026).

### 7. Prediction

The performance of first- and second-order, Lasso- and Ridge-regularized linear models was evaluated via repeated random split cross-validation across a range of training set sizes.

Training fractions ranged from 0.05 to 0.95 in intervals of 0.05. For each fraction, we trained the models on 100 different training sets, each an independent random sample from the full dataset. The remaining observations were used as the test set. For each split, models were fit using the cv.glmnet function from the glmnet R package (Friedman et al., 2010), with the regularization parameter λ selected via inner 10-fold cross-validation on the training set, using the value minimizing cross-validated error. Out-of-sample performance was then assessed by generating predictions on the test set and computing *R*^2^, using only the test set observations in both the residual sum of squares and the total sum of squares (see Eq. 4 in the main text).

## Notes

### Competing Interest Statement

The authors have declared no competing interest.

