## Supplementary Information (Suppl. Figures and Table from Statistical learning of bacterial growth in constructed environments). for "Statistical learning of bacterial growth in combinatorially constructed environments"

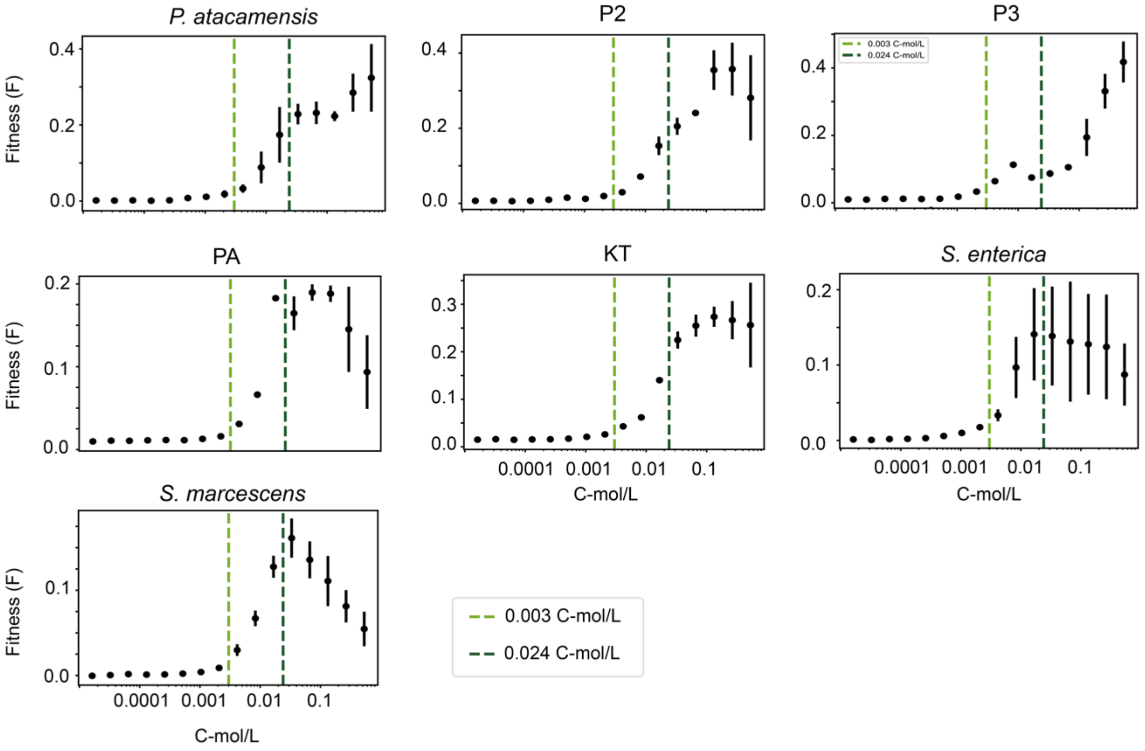


***Figure S1. Fitness after 24 hours of growth in M9 supplemented with a glucose concentration gradient.*** *To identify the carbon concentration range at which growth remains carbon-limited, seven bacterial strains (*Pseudomonas atacamensis*, P2, P3, PA, KT,* Salmonella enterica*, and* Serratia marcescens*) were grown in M9 minimal medium (base medium) supplemented with increasing glucose concentrations (0.0001–0.1 C-mol/L). Points represent biological replicate measurements and black error bars indicate mean ± standard deviation. Light green and dark green dashed lines indicate the carbon molarity in media when a single resource or all eight resources are added, respectively.*

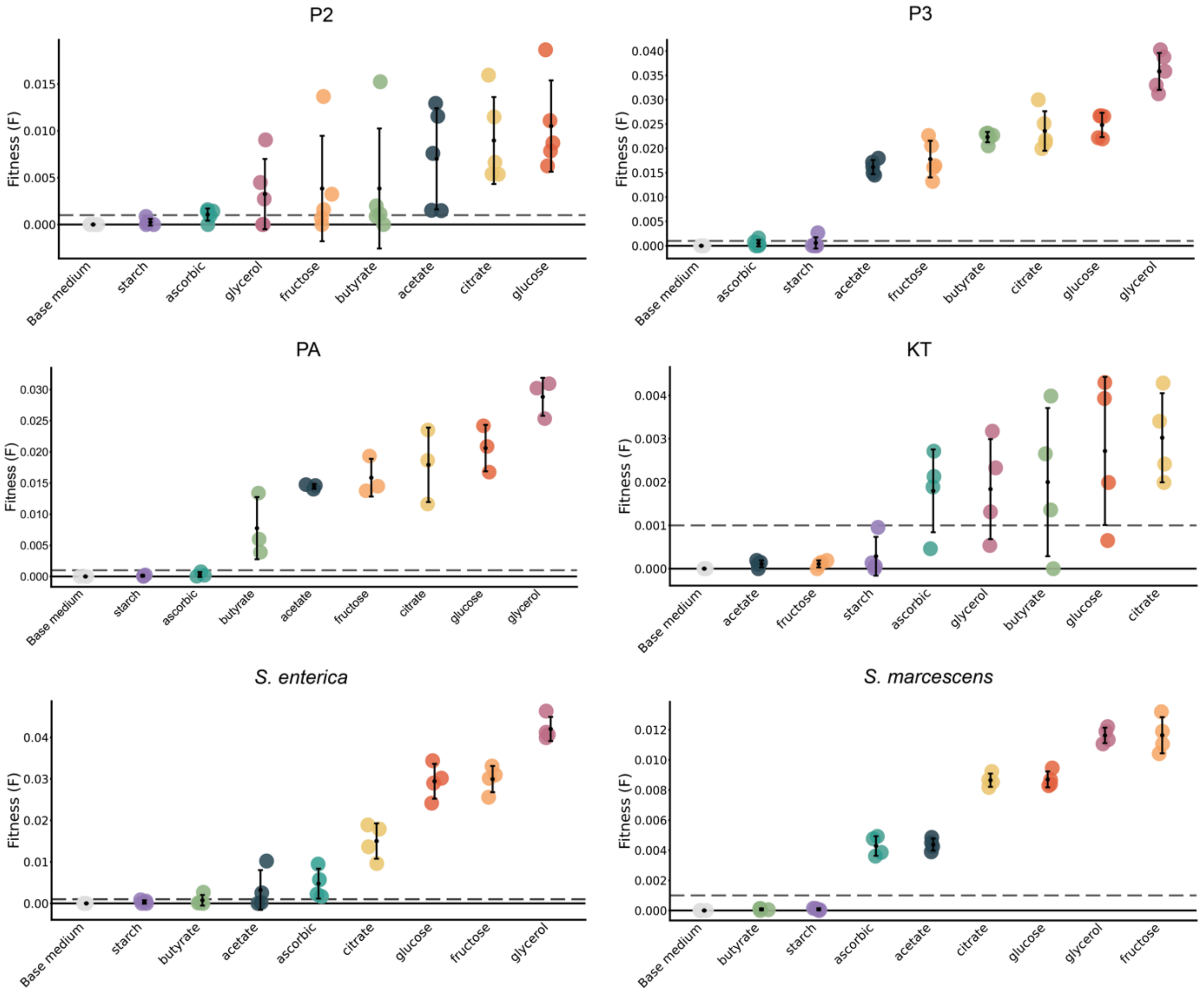


***Figure S2. Individual fitness on single carbon sources***. *Fitness value of each strain when grown on base medium supplemented exclusively with each of the eight carbon sources tested. Error bars represent standard deviation across replicates. The dashed horizontal line indicates the detection limit (0.001). Substantial variation in strain-specific utilization efficiency is evident across carbon sources.*

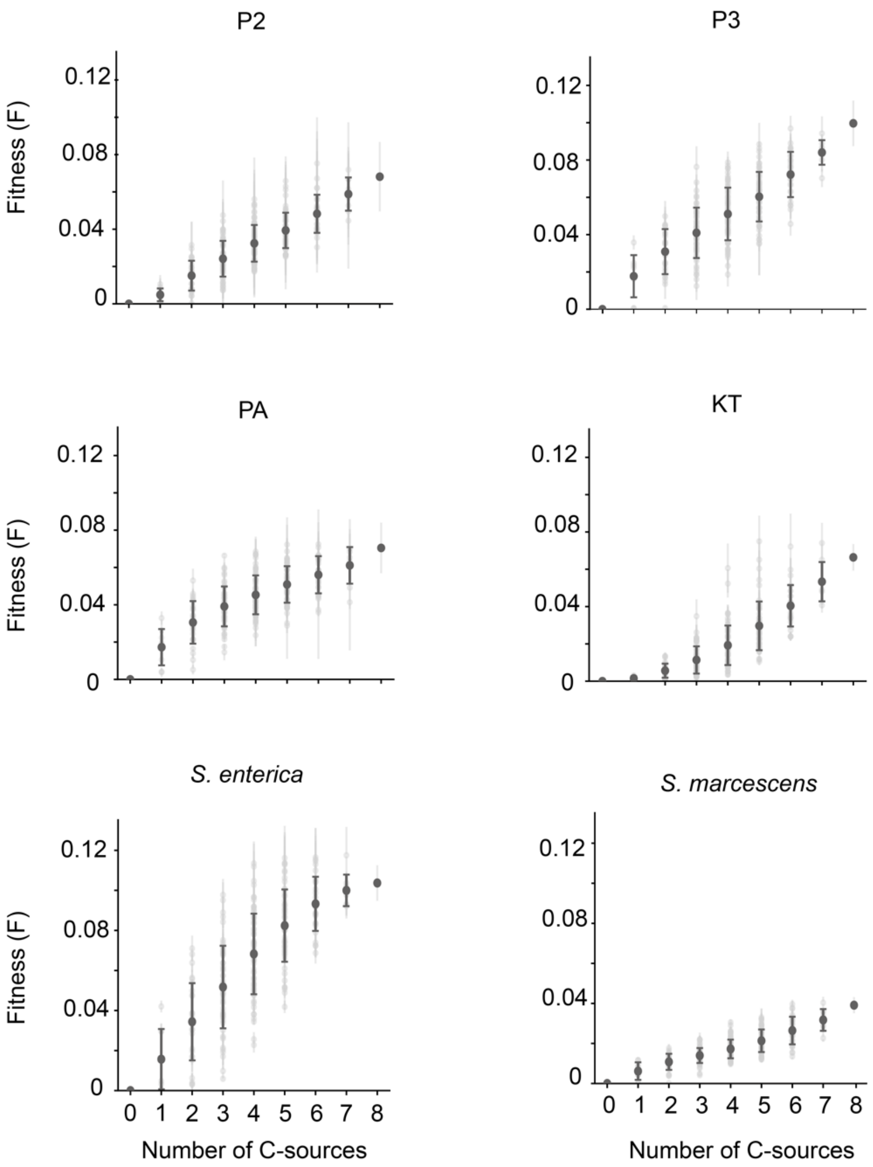


***Figure S3. Resource diversity and fitness relationships across strains.*** *Fitness (*F*) measured across 255 environments plotted against the number of carbon sources in the mixture. Light grey points represent individual fitness of each strain in each individual environment; dark grey points indicate the average fitness (± standard deviation) as a function of the number of carbon sources present in the mixture.*

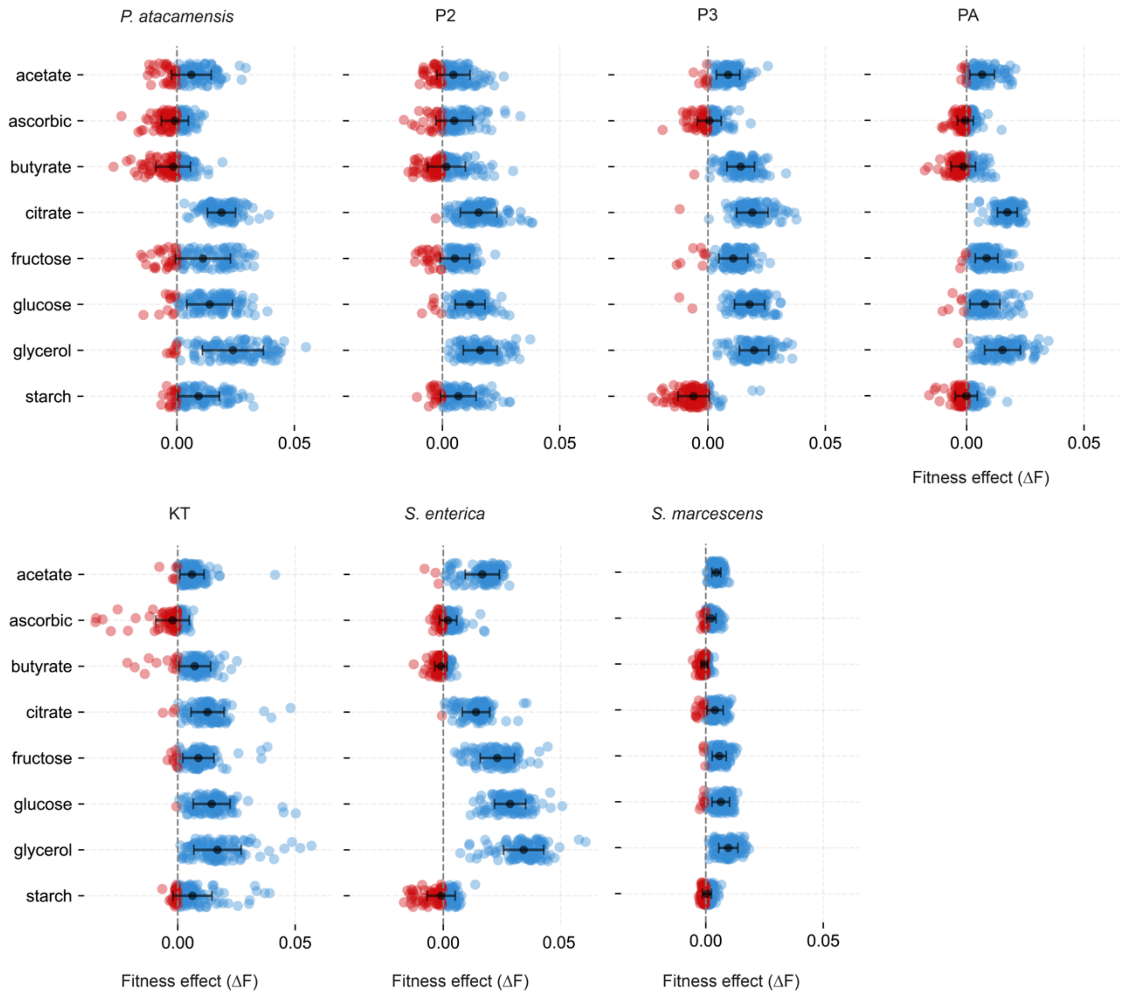


***Figure S4. Context-dependent dual behavior of carbon sources across strains.*** *Fitness effects distribution for each carbon source (given on the y-axis) added to every possible background environment across strains. Blue and red dots indicate environments in which the resource acts as a fitness enhancer or fitness inhibitor, respectively.*

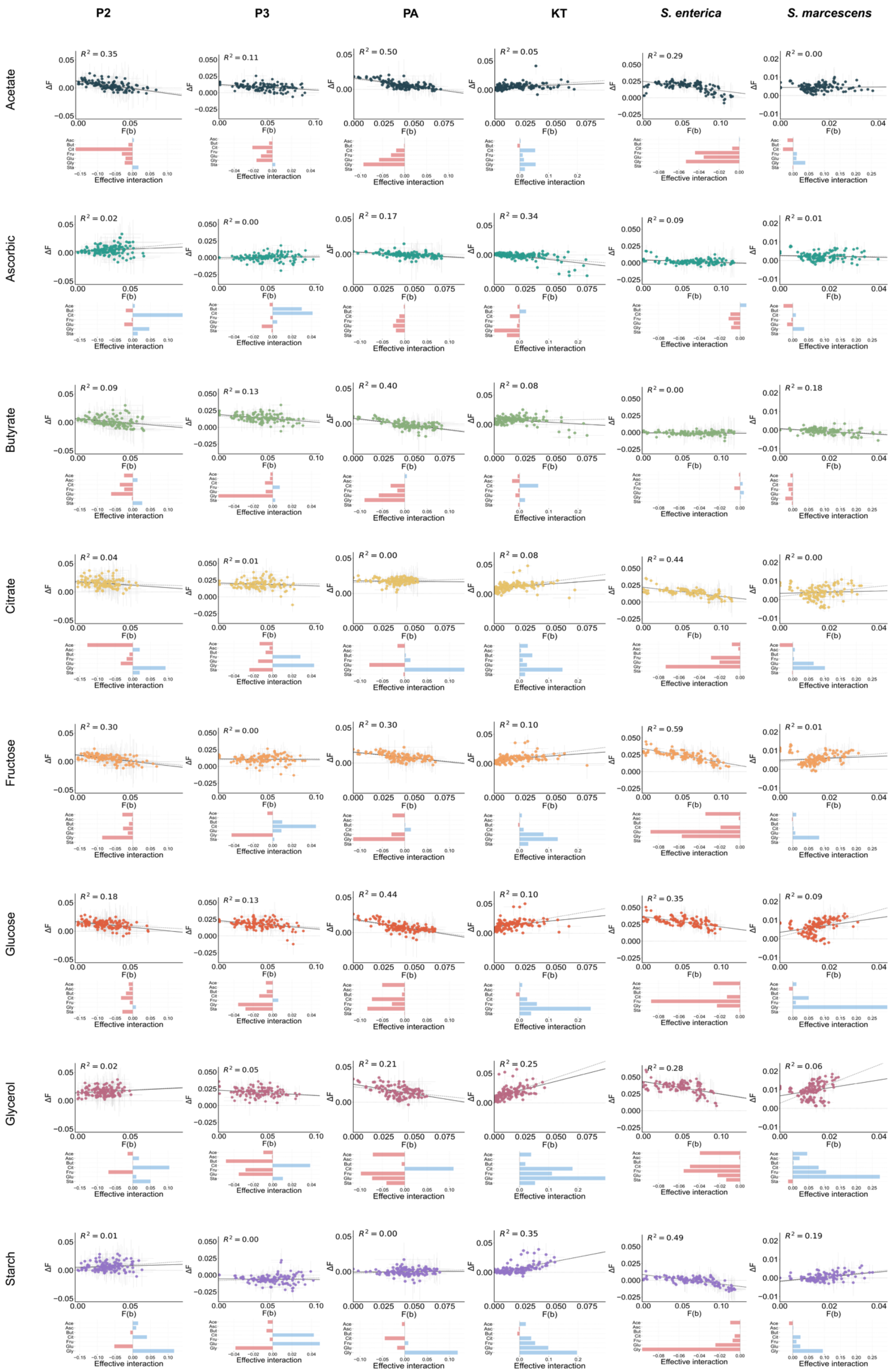

P2
P3
PA
KT
S. enterica
S. marcescens
Acetate
R² = 0.35
R² = 0.11
R² = 0.50
R² = 0.05
R² = 0.29
R² = 0.00
Ascorbic
R² = 0.02
R² = 0.00
R² = 0.17
R² = 0.34
R² = 0.09
R² = 0.01
Butyrate
R² = 0.09
R² = 0.13
R² = 0.40
R² = 0.08
R² = 0.00
R² = 0.18
Citrate
R² = 0.04
R² = 0.01
R² = 0.00
R² = 0.08
R² = 0.44
R² = 0.00
Fructose
R² = 0.30
R² = 0.00
R² = 0.30
R² = 0.10
R² = 0.59
R² = 0.01
Glucose
R² = 0.18
R² = 0.13
R² = 0.44
R² = 0.10
R² = 0.35
R² = 0.09
Glycerol
R² = 0.02
R² = 0.05
R² = 0.21
R² = 0.25
R² = 0.28
R² = 0.06
Starch
R² = 0.01
R² = 0.00
R² = 0.00
R² = 0.35
R² = 0.49
R² = 0.19
ΔF
F(b)
Effective interaction

***Figure S5. Fitness effects across strains.*** *Fitness effects of adding a carbon source (color-coded and positioned on the y-axis) to a background environment are well predicted by linear models linking fitness effects to the fitness of the background environment. Panels show results for P2, P3, PA, KT,* S. enterica *and* S. marcescens *(indicated at the bottom). Data represent mean ± standard deviation across replicates.*

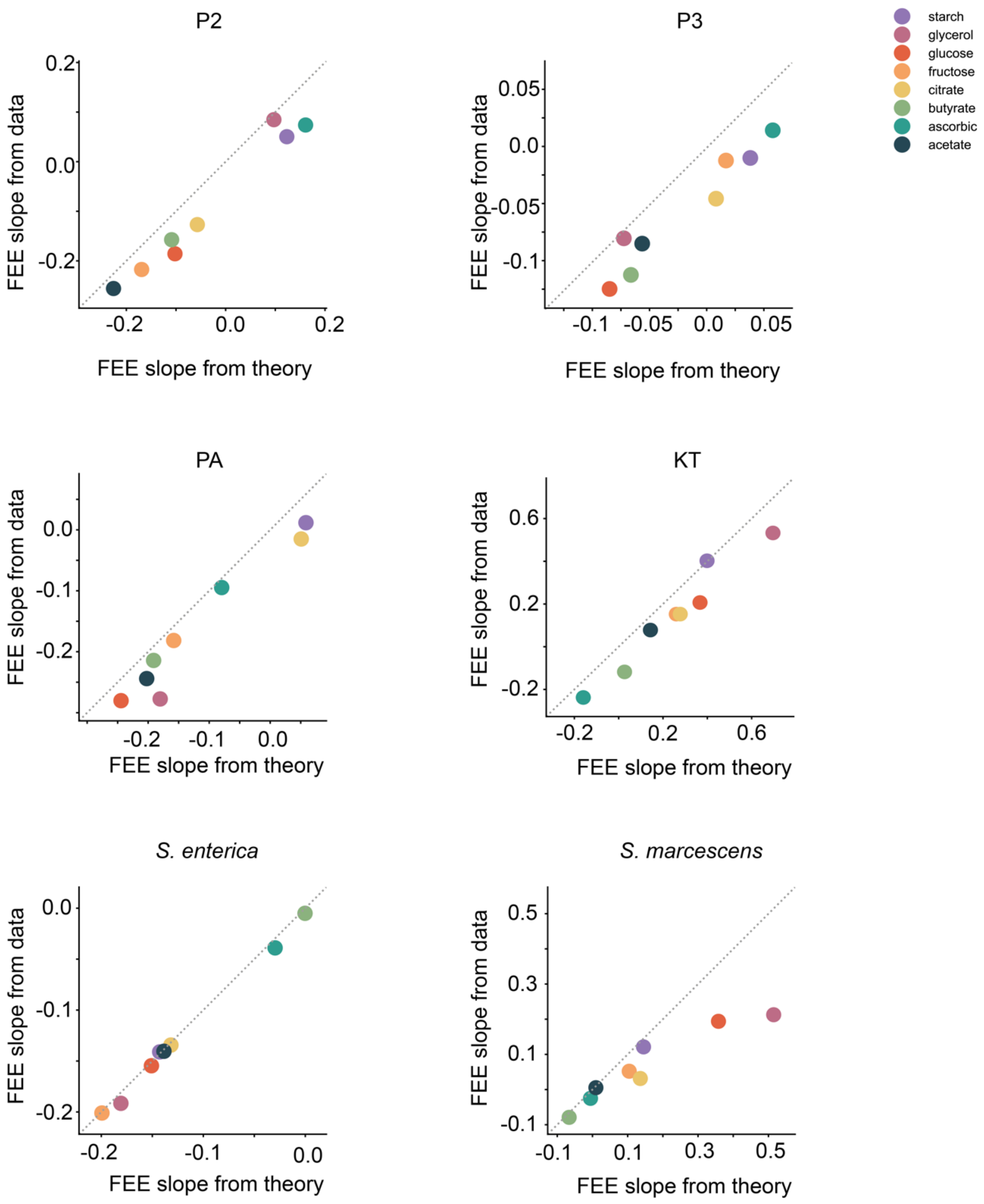


***Figure S6. Empirical vs theoretical FEE slopes.*** *FEE slopes from data are compared with theoretical predictions across strains.*

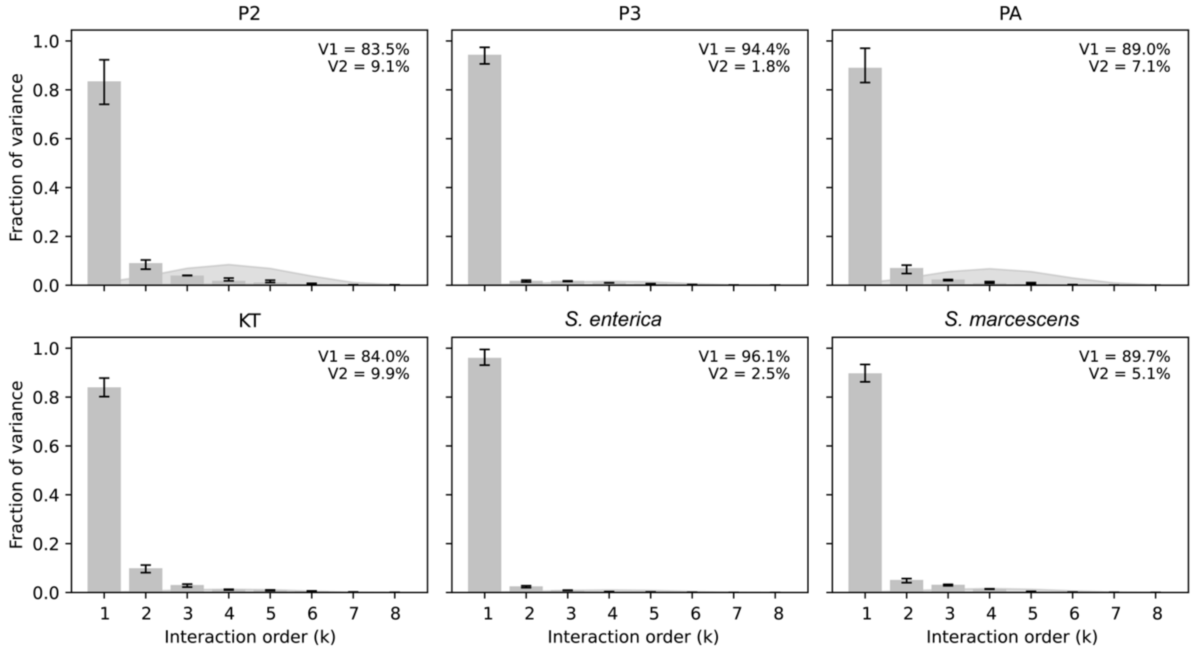


***Figure S7. Contribution of interactions at all orders to fitness variation across strains.*** *Fitness variance is predominantly explained by additive and pairwise effects across all environments across strains. Consistent with the findings for* P. atacamensis *(Figure 3C), the results are quantitatively and qualitatively similar, demonstrating that higher-than-pairwise interactions have a negligible effect on the fitness impact of new resources under these conditions.*

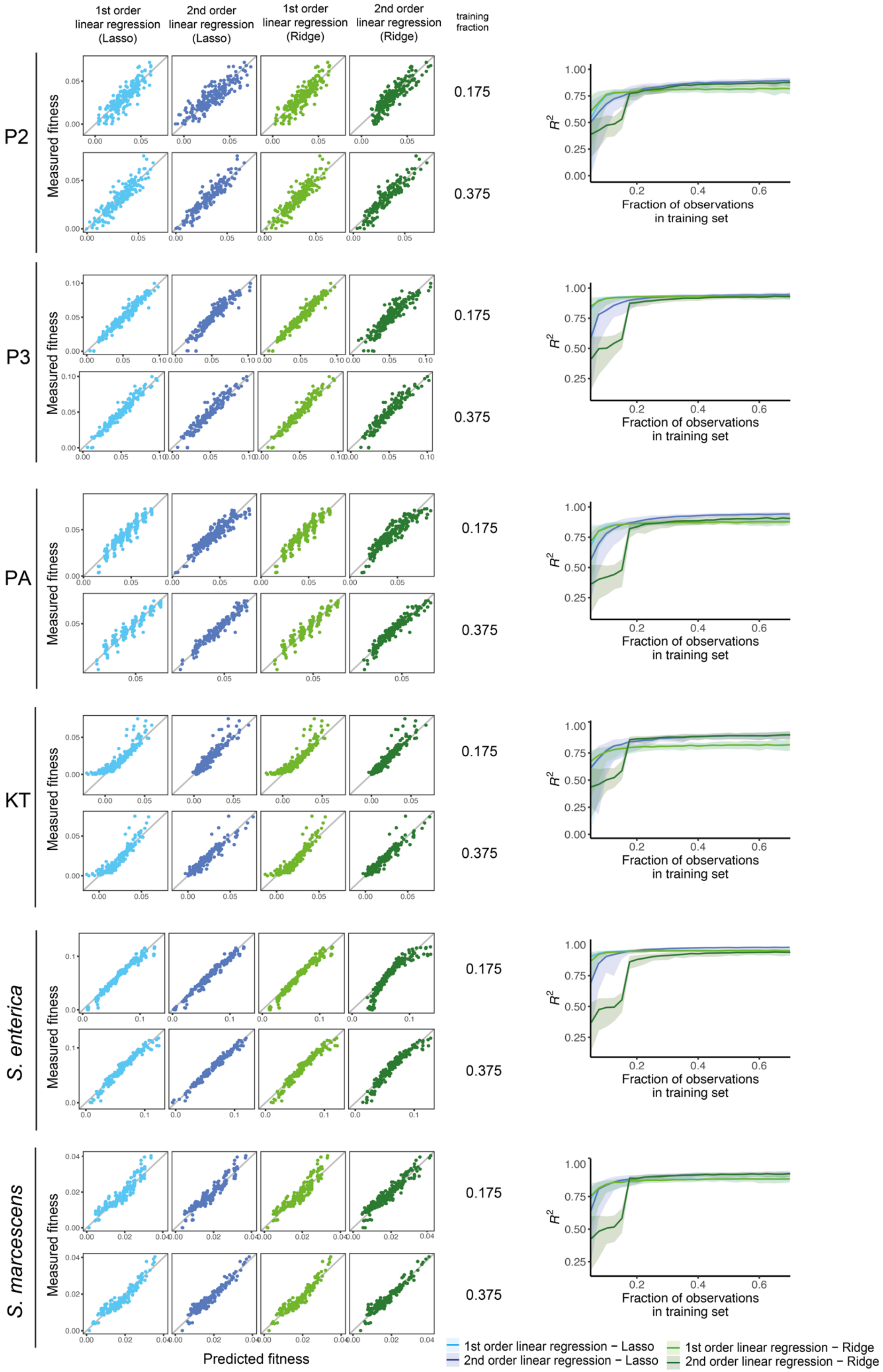

1st order linear regression (Lasso)
2nd order linear regression (Lasso)
1st order linear regression (Ridge)
2nd order linear regression (Ridge)
training fraction
P2
P3
PA
KT
S. enterica
S. marcescens
Measured fitness
Predicted fitness
0.175
0.375
$R^2$
Fraction of observations in training set
1st order linear regression − Lasso
2nd order linear regression − Lasso
1st order linear regression − Ridge
2nd order linear regression − Ridge

***Figure S8. Model prediction accuracy across bacterial isolates and species.*** *(Left panels) Measured versus predicted fitness values for first-order and second-order linear regression models (Lasso and Ridge) at training set fractions of 0.175 and 0.375 across strains. (Right panels) Relationship between training set fraction and prediction accuracy ($R^2$) with shaded interquartile ranges. Second-order models (dark colors) consistently outperform first-order models (light colors) across all species. Both Lasso and Ridge regularization show similar performance with Ridge achieving slightly higher $R^2$ values at higher training fractions.*

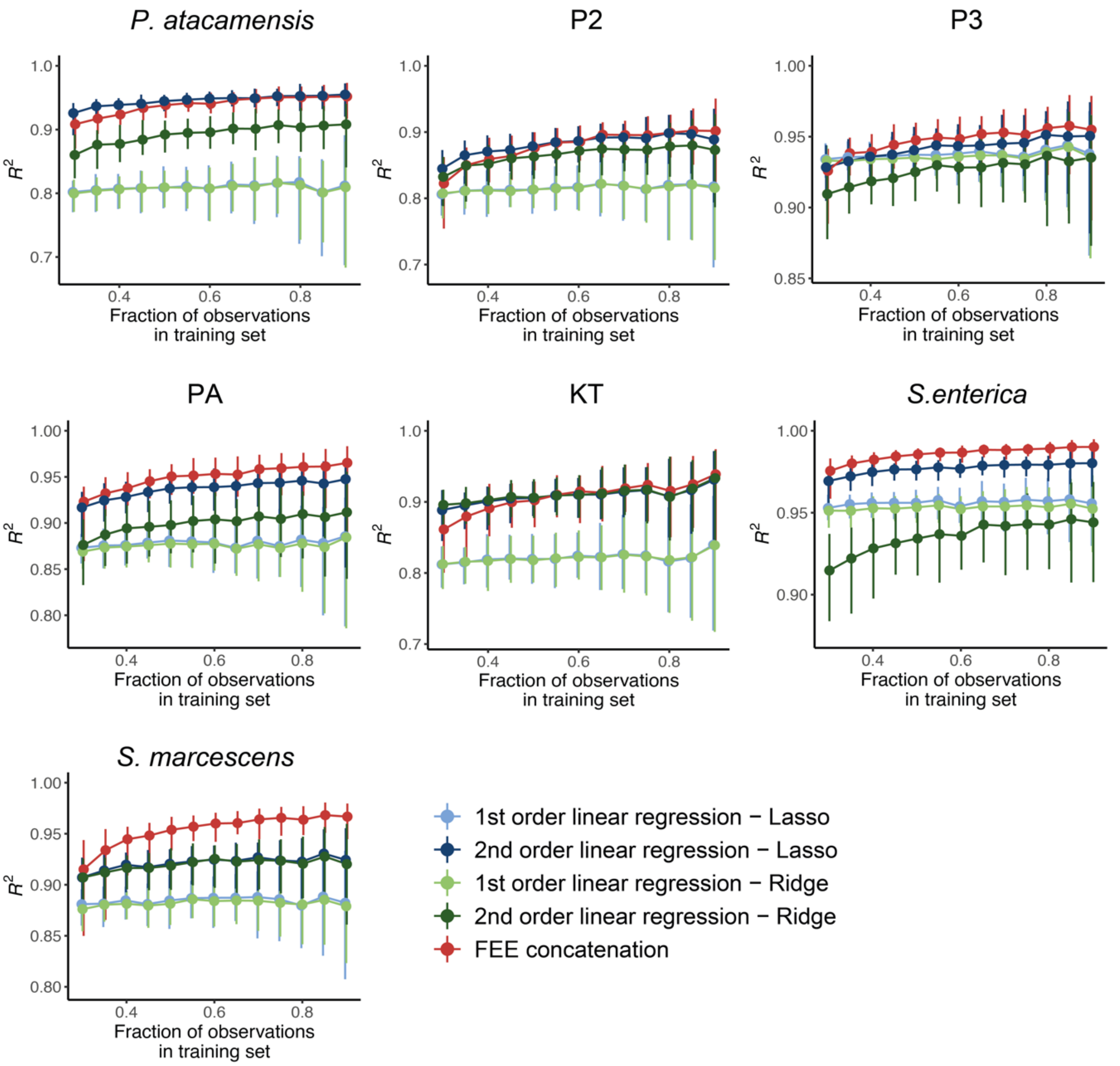


***Figure S9. Linear regression models and FEE concatenation method accurately predict microbial growth out of sample across bacterial species.*** *Relationship between training set fraction and model prediction accuracy ($R^2$) across strains. Each panel shows median $R^2$ values across 100 independent random partitions with shaded error bars representing the interquartile range. Models compared include first-order and second-order linear regression with Lasso (light and dark blue, respectively) and Ridge regularization (light and dark green, respectively), as well as an alternative FEE concatenation method (red). Results are qualitatively and quantitatively similar across all species, demonstrating the robustness and generalizability of the predictive modeling approach. Second-order models consistently outperform first-order models across all species and training fractions, while FEE concatenation achieves comparable or superior performance to linear regression, particularly at higher training set fractions.*

### Supplementary material

*Table 1: Composition and concentrations of carbon sources used in the study, including their chemical nature, supplier information, and reference identifiers.*

<table>
<tr><th>Reagent type</th><th>Carbon source (CS)</th><th>C-mol/L</th><th>1X</th><th>C-mol/ L (8X)</th><th>8X</th><th>Supplier</th><th>Reference</th></tr>
<tr><td>Chemical compound, drug</td><td>Sodium acetate anhydrous</td><td rowspan="8">0.003</td><td>0.003 C-mol/L</td><td rowspan="8">0.024</td><td>0.024 C-mol/L</td><td>Sigma Aldrich</td><td>W302406</td></tr>
<tr><td>Chemical compound, drug</td><td>L-ascorbic acid</td><td>0.088 g/L</td><td>0.7 g/L</td><td>Sigma Aldrich</td><td>A92902</td></tr>
<tr><td>Chemical compound, drug</td><td>Sodium butyrate</td><td>0.083 g/L</td><td>0.664 g/L</td><td>Sigma Aldrich</td><td>B5887</td></tr>
<tr><td>Chemical compound, drug</td><td>Sodium citrate monobasic</td><td>0.003 C-mol/L</td><td>0.024 C-mol/L</td><td>Sigma Aldrich</td><td>71498</td></tr>
<tr><td>Chemical compound, drug</td><td>D-(-)-Fructose</td><td>0.009%</td><td>0.072%</td><td>Sigma Aldrich</td><td>F0127</td></tr>
<tr><td>Chemical compound, drug</td><td>D-(+)-Glucose</td><td>0.009%</td><td>0.072%</td><td>Sigma Aldrich</td><td>G7021</td></tr>
<tr><td>Chemical compound, drug</td><td>Glycerol</td><td>0.009%</td><td>0.072%</td><td>Sigma Aldrich</td><td>G9012</td></tr>
<tr><td>Chemical compound, drug</td><td>Starch, soluble</td><td>0.002%</td><td>0.016%</td><td>Sigma Aldrich</td><td>S9765</td></tr>
</table>
